## Supplementary Materials for "Fitness estimation for viral variants in the context of cellular coinfection"

### Supplementary Materials to the Manuscript: Fitness estimation for viral variants in the context of cellular coinfection

Huisheng Zhu[*,1], Brent E Allman[*,2], Katia Koelle[1,3,†]

[*]Equal contributions

[1]Department of Biology, Emory University, Atlanta, Georgia 30322

[2]Graduate Program in Population Biology, Ecology, and Evolution, Emory University, Atlanta, Georgia 30322

[3]Emory-UGA Center of Excellence for Influenza Research and Surveillance (CEIRS), Atlanta, GA, USA

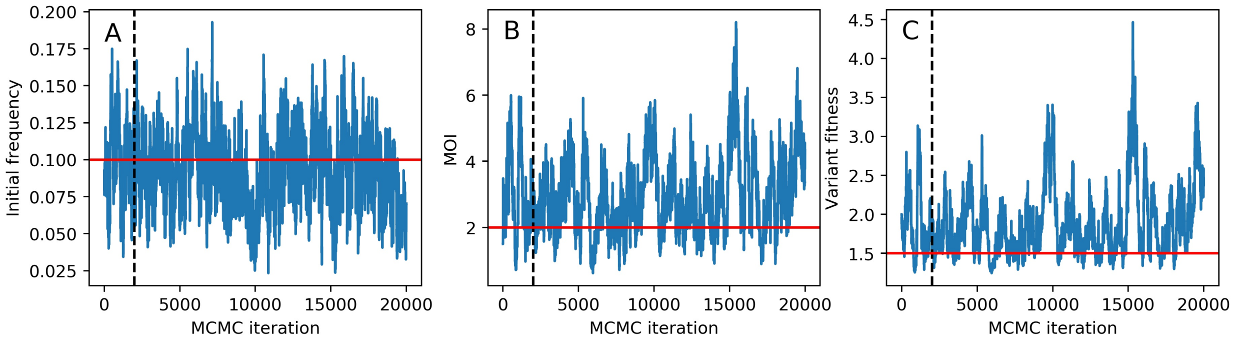


Figure S1: MCMC trace plots for parameters estimated by interfacing the deterministic within-host model with the simulated data. (A) Trace plot for initial frequency of the variant. (B) Trace plot for mean cellular multiplicity of infection. (C) Trace plot for variant fitness. 20,000 MCMC iterations were run. Following the removal of the first 2,000 MCMC iterations as burn-in, the MCMC chain was sampled every 50 iterations.

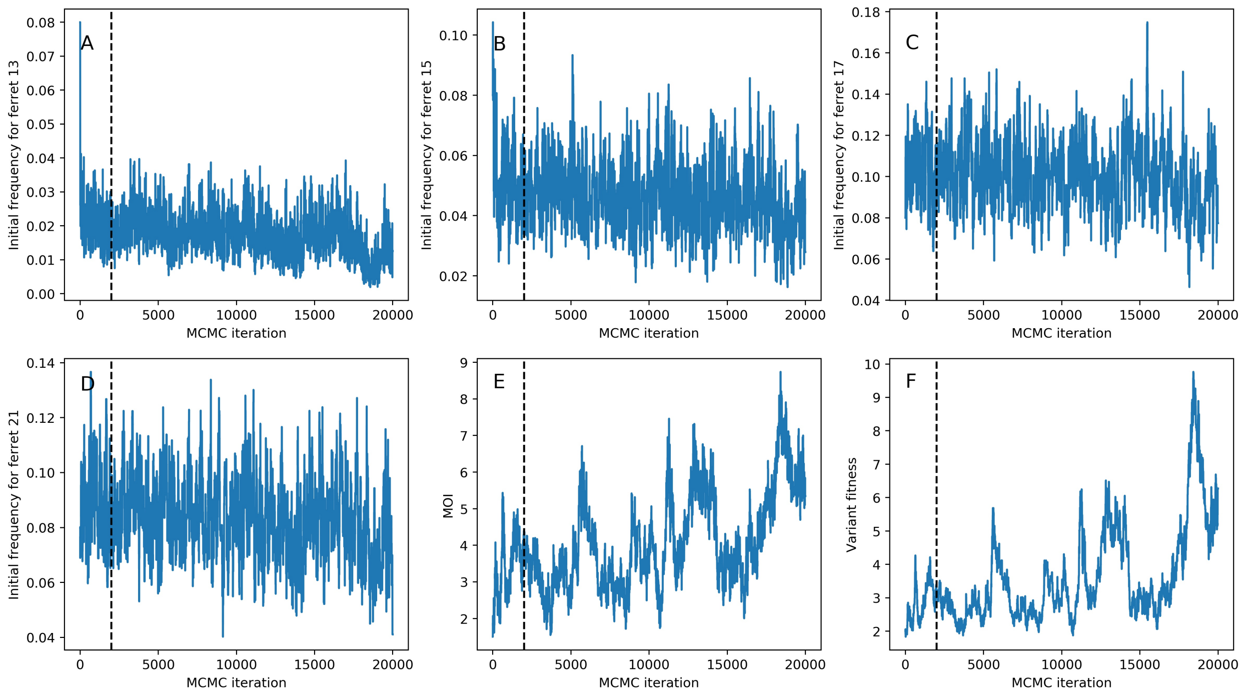


Figure S2: MCMC trace plots for parameters estimated by interfacing the deterministic within-host model with the influenza H5N1 experimental challenge study data. (A) Trace plot for initial frequency of the G788A variant in ferret 13. (B) Trace plot for initial frequency of the G788A variant in ferret 15. (C) Trace plot for initial frequency of the G788A variant in ferret 17. (D) Trace plot for initial frequency of the G788A variant in ferret 21. (E) Trace plot for mean cellular multiplicity of infection. (F) Trace plot for G788A variant fitness. 20,000 MCMC iterations were run. Following the removal of the first 2,000 MCMC iterations as burn-in, the MCMC chain was sampled every 50 iterations.

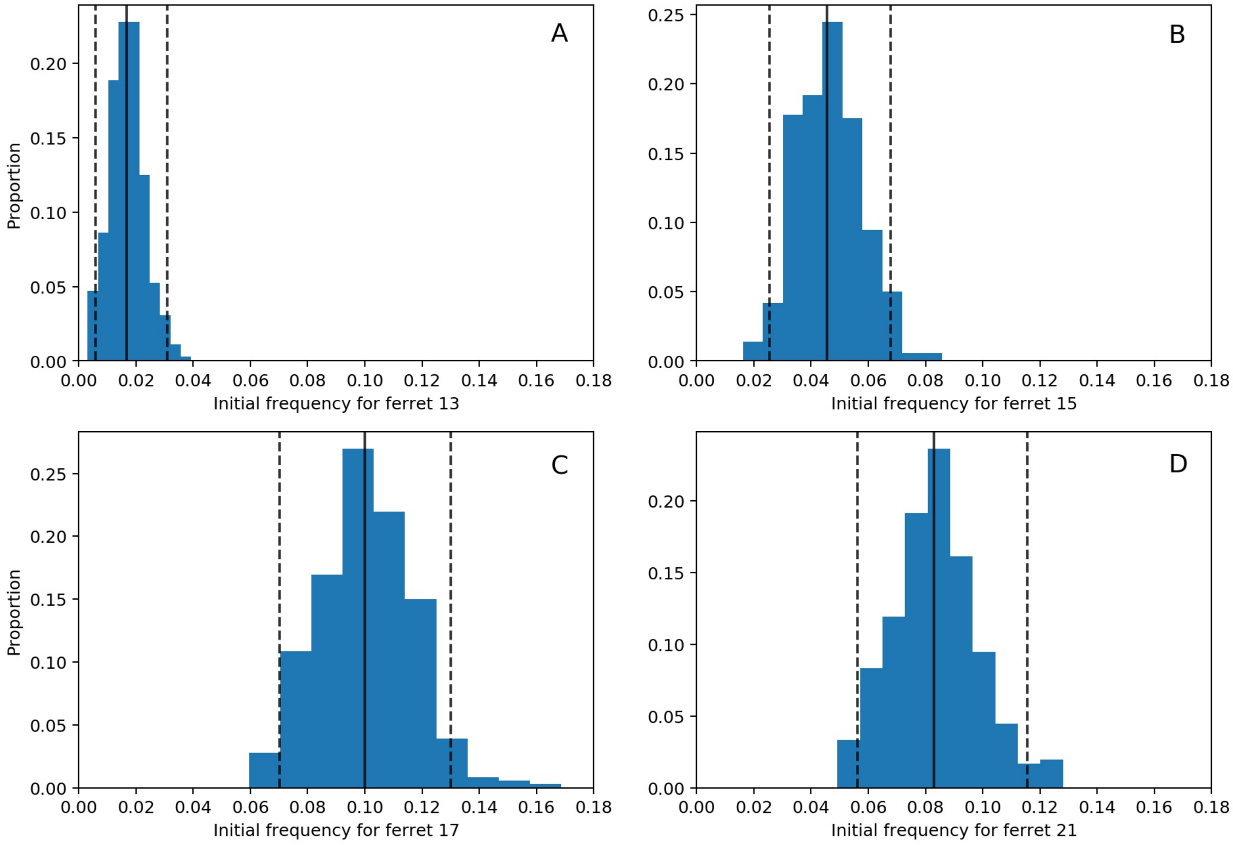


Figure S3: Posterior distributions of initial G788A frequencies for (A) ferret 13, (B) ferret 15, (C) ferret 17, and (D) ferret 21, from fitting the deterministic within-host model. In (A)-(D), black solid lines show the median values of the posterior densities and black dashed lines show the 95% credible intervals.

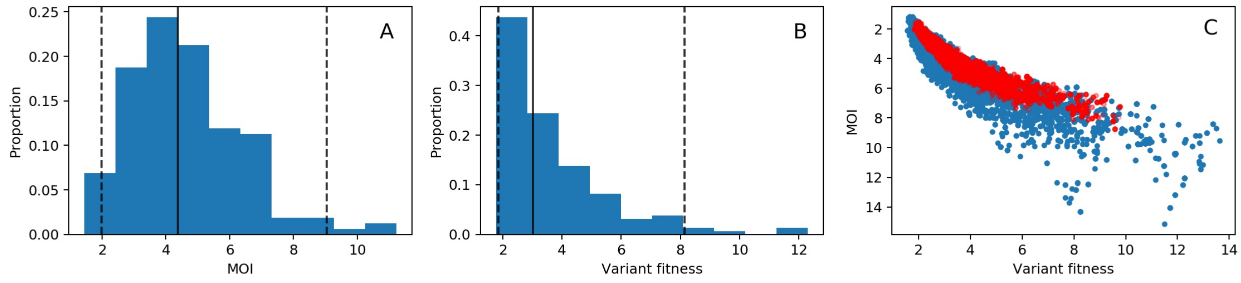


Figure S4: Parameter estimation for variant G788A, assuming deterministic within-host dynamics and measurement noise of $\nu = 25$. (A) Posterior distribution for the mean cellular multiplicity of infection. (B) Posterior distribution for variant fitness. In (A) and (B), black solid lines show the median values of the posterior densities and black dashed lines show the 95% credible intervals. (C) Joint density plot for MOI and variant fitness (blue). For comparison, we have superimposed the joint density plot shown in Figure 3D (red).

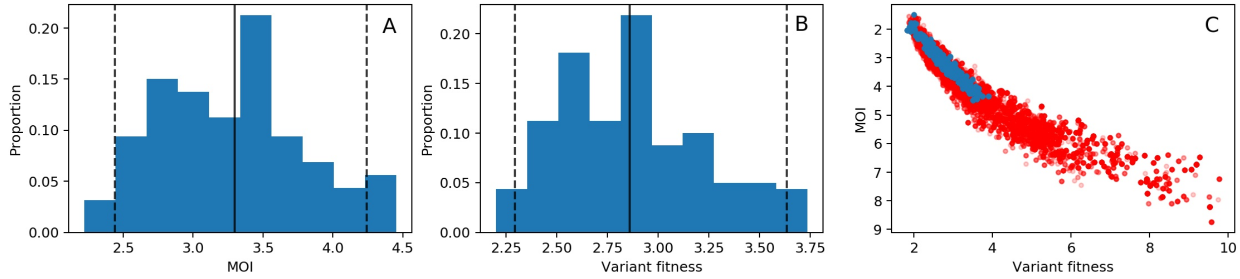


Figure S5: Parameter estimation for variant G788A, assuming deterministic within-host dynamics and measurement noise of $\nu = 400$. (A) Posterior distribution for the mean cellular multiplicity of infection. (B) Posterior distribution for variant fitness. In (A) and (B), black solid lines show the median values of the posterior densities and black dashed lines show the 95% credible intervals. (C) Joint density plot for MOI and variant fitness (blue). For comparison, we have superimposed the joint density plot shown in Figure 3D (red).

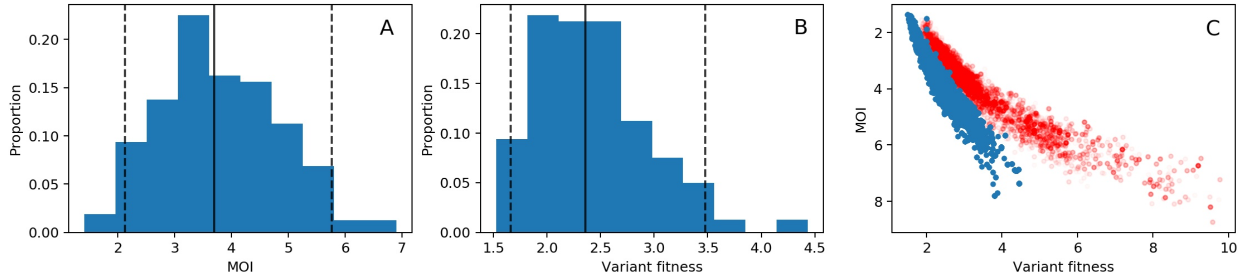


Figure S6: Parameter estimation for variant G788A, assuming deterministic within-host dynamics and a viral generation time of 6 hours. (A) Posterior distribution for the mean cellular multiplicity of infection. (B) Posterior distribution for variant fitness. In (A) and (B), black solid lines show the median values of the posterior densities and black dashed lines show the 95% credible intervals. (C) Joint density plot for MOI and variant fitness (blue). For comparison, we have superimposed the joint density plot shown in Figure 3D (red).

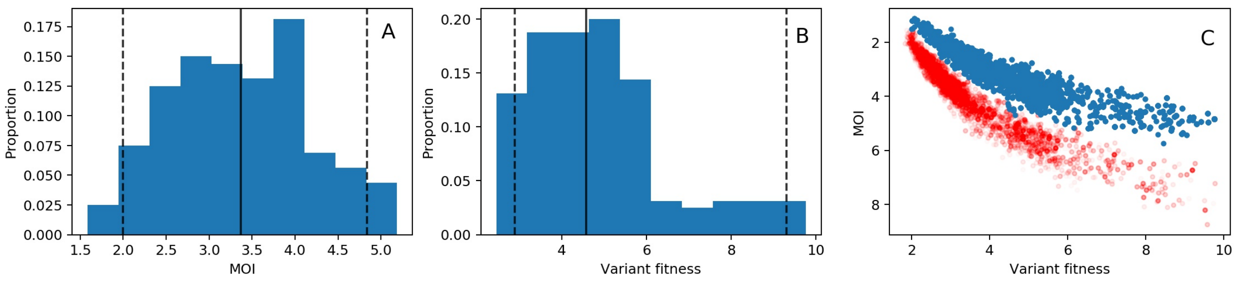


Figure S7: Parameter estimation for variant G788A, assuming deterministic within-host dynamics and a viral generation time of 12 hours. (A) Posterior distribution for the mean cellular multiplicity of infection. (B) Posterior distribution for variant fitness. In (A) and (B), black solid lines show the median values of the posterior densities and black dashed lines show the 95% credible intervals. (C) Joint density plot for MOI and variant fitness (blue). For comparison, we have superimposed the joint density plot shown in Figure 3D (red).

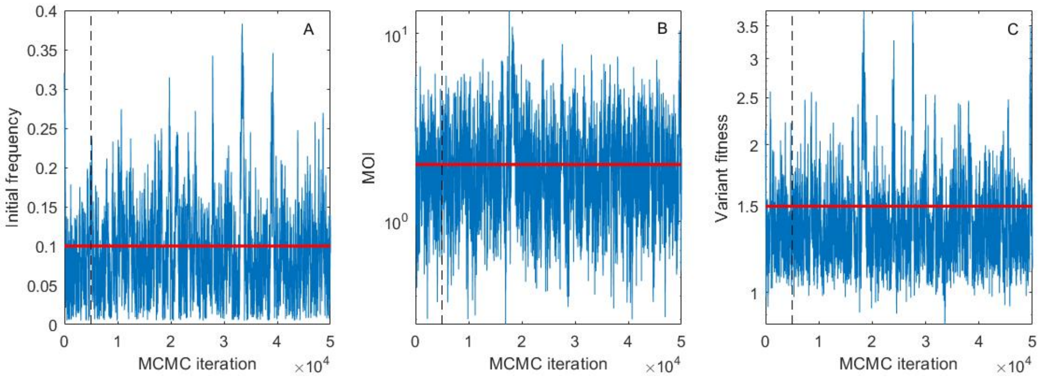


Figure S8: MCMC trace plots for parameters estimated by interfacing the stochastic within-host model with the simulated data. (A) Trace plot for initial frequency of the variant. (B) Trace plot for mean cellular multiplicity of infection. (C) Trace plot for variant fitness. 50,000 MCMC iterations were run. Following the removal of the first 5,000 MCMC iterations as burn-in, the MCMC chain was sampled every 100 iterations.

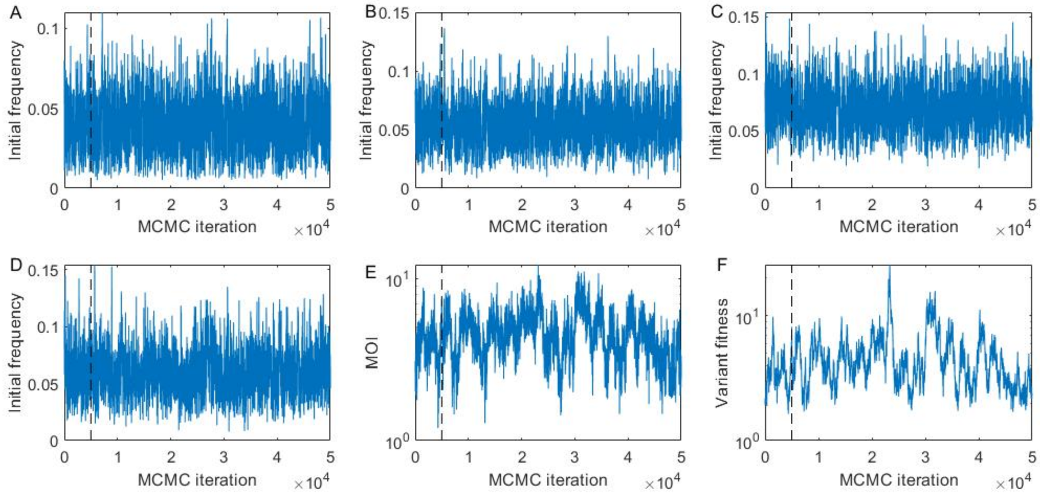


Figure S9: MCMC trace plots for parameters estimated by interfacing the stochastic within-host model with the influenza H5N1 experimental challenge study data. (A) Trace plot for initial frequency of the G788A variant in ferret 13. (B) Trace plot for initial frequency of the G788A variant in ferret 15. (C) Trace plot for initial frequency of the G788A variant in ferret 17. (D) Trace plot for initial frequency of the G788A variant in ferret 21. (E) Trace plot for mean cellular multiplicity of infection. (F) Trace plot for G788A variant fitness. 50,000 MCMC iterations were run. Following the removal of the first 5,000 MCMC iterations as burn-in, the MCMC chain was sampled every 100 iterations.

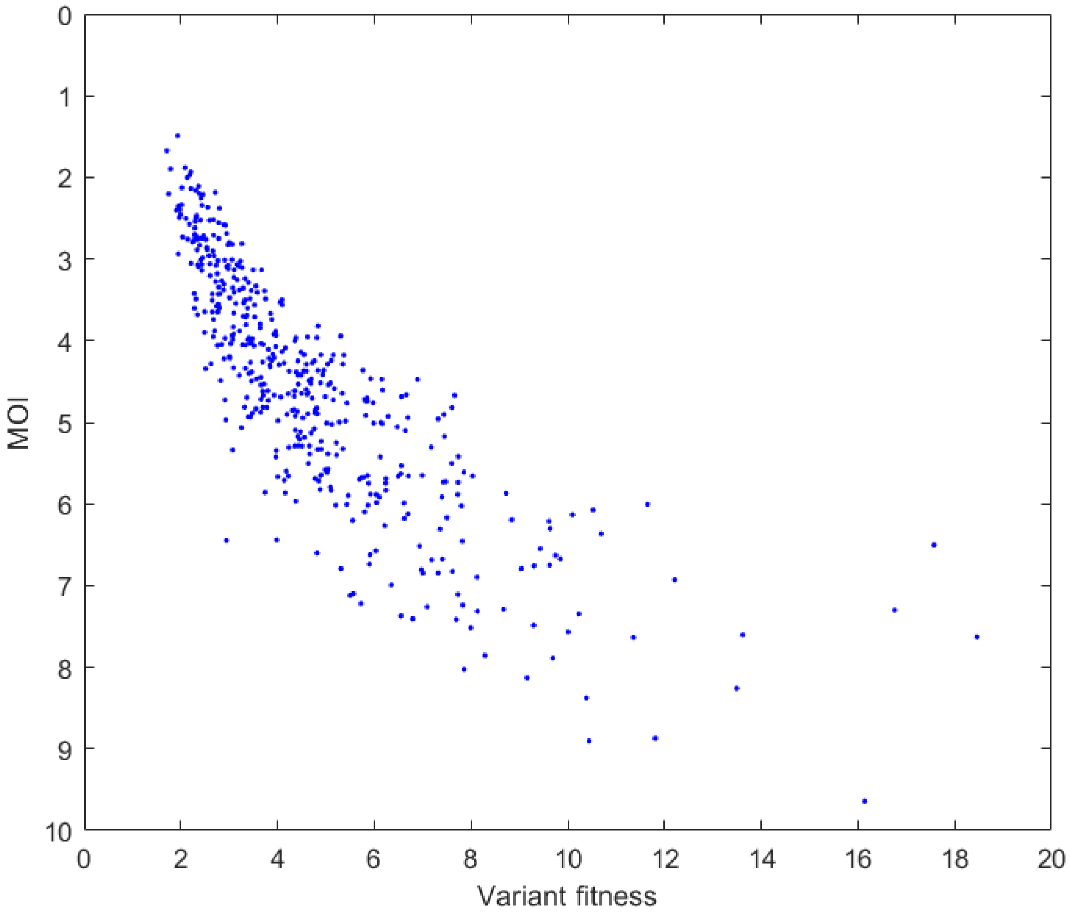


Figure S10: Joint density of MOI versus variant fitness for the stochastic model fit to the influenza H5N1 experimental challenge study data.

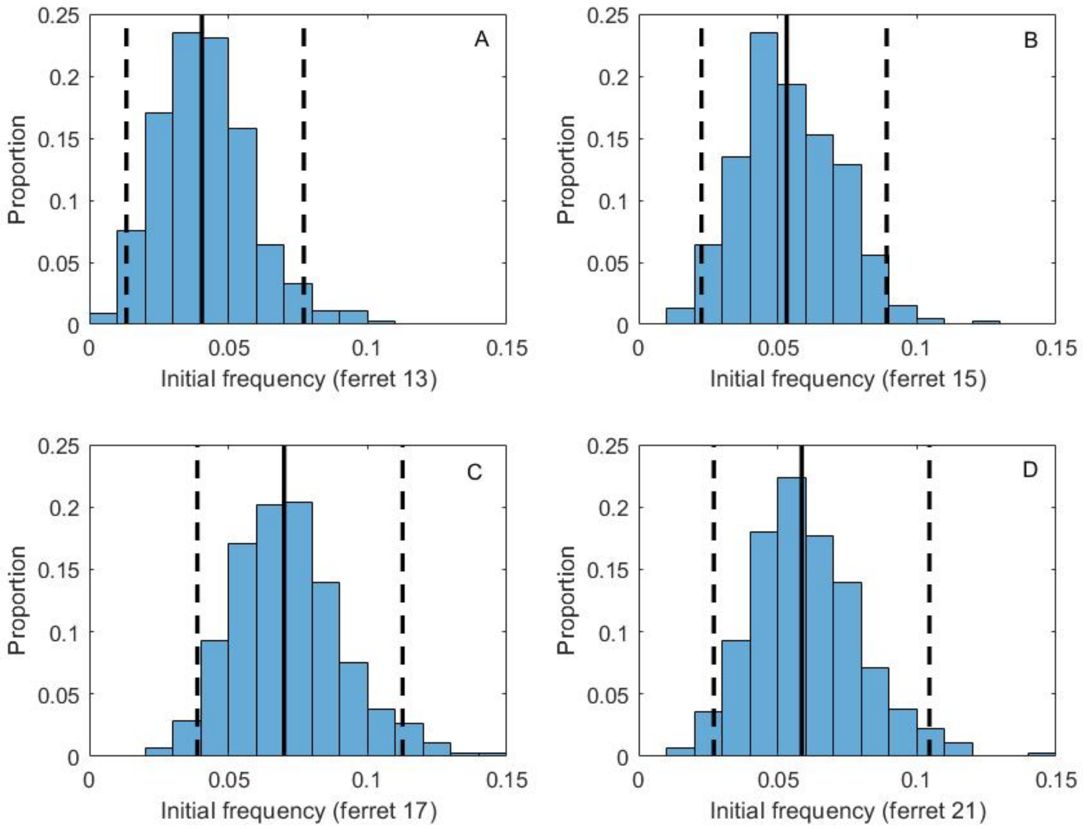


Figure S11: Posterior distributions of initial G788A frequencies for (A) ferret 13, (B) ferret 15, (C) ferret 17, and (D) ferret 21, from fitting the stochastic within-host model. In (A)-(D), black solid lines show the median values of the posterior densities and black dashed lines show the 95% credible intervals.

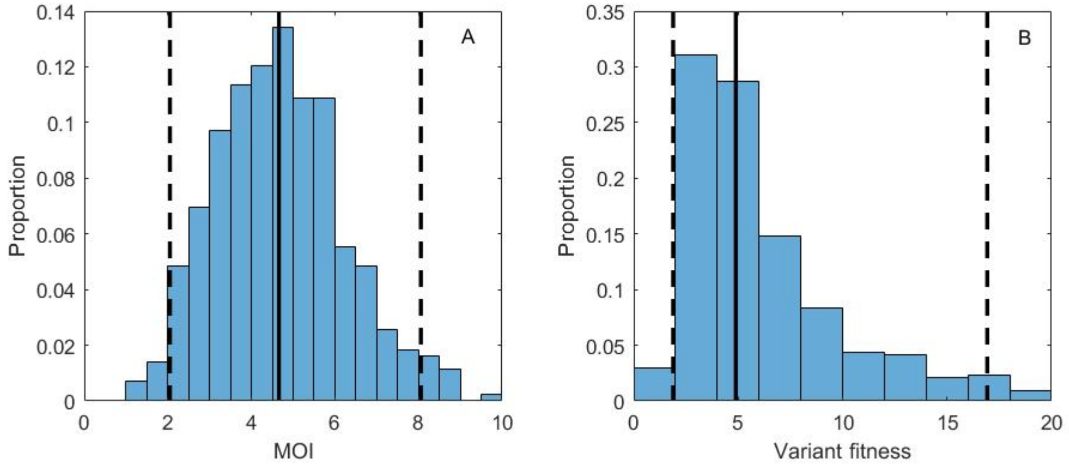


Figure S12: Fitness estimation for variant G788A, assuming stochastic within-host dynamics, parameterized with an effective viral population size of $N = 40$. (A) Posterior distribution for the mean cellular multiplicity of infection $M$. (B) Posterior distribution for variant fitness. In (A) and (B) black solid lines show the median values of the posterior densities and black dashed lines show the 95% credible intervals.
